# “Tracking the Rain Bird”: Modeling the monthly distribution of Pied cuckoo in India

**DOI:** 10.1101/2020.11.06.371674

**Authors:** Debanjan Sarkar, Bharti Tomar, R. Suresh Kumar, Sameer Saran, Gautam Talukdar

## Abstract

Pied cuckoo *Clamator jacobinus* (Boddart, 1783) is a migratory, brood-parasitic bird found in the African and Indian Subcontinent. Although the southern Indian population is presumably resident, the North Indian Population migrates from Africa to India during the summer. The arrival of the bird is linked to the onset of monsoon in India from scientific literature to folklore. It is known to make its appearance in central and northern India in the last week of May or early June, indicating the imminent arrival of the monsoon with its unmistakably loud metallic calls. There have been few attempts to compile relevant information on the species migration in the early 1900s and citizen science approach by Bird-count India; little information is available on how environmental factors might be affecting its migration. Here, we have used Maximum Entropy modeling to identify the monthly and seasonal distribution patterns and major bioclimatic factors that might be influencing the distribution of the species in India. We have used E-Bird citizen science platform data, seven bioclimatic variables, and monthly NDVI of respective months for building the models. The predicted output shows the species presence throughout the year in southern India. In contrast, in northern India, the distribution is dynamic, peaking in summers in the Month of May-June and no presence in winter. The influence of bioclimatic variables used in SDM varied monthly; Water vapor pressure was the primary contributing variable in the months prior to species arrival. In July, it was NDVI (Higher NDVI suggests abundance of food resources for the species). In August-September, Windspeed and water vapor pressure (Factors might be responsible for the departure of the species) have contributed highest. Our approach provides a more concise understanding of Pied cuckoo’s monthly distributions throughout India, which helps understand the complex seasonal shifts in the distribution of such migratory birds.

## 1. Introduction

Birds have been the subject of theoretical developments in behaviour, ecology, and evolutionary biology (Hackett et al., 2008; Schluter, 1996; Sutherland et al., 2004;), and yet their distribution and migration patterns have always been a question with varying hypothesis amidst the ornithological community. Seasonal resource availability results in the varying seasonal distributions of many birds (Engler et al. 2014; Eyres et al. 2017) and seasonal niches in time and space considerations on ecological niches are more complex in organisms with seasonal distributions (Engler et al. 2014; Martínez-Meyer et al. 2004; Nakazawa et al. 2004,). Typical migratory birds demonstrate a breeding and a non-breeding distribution. Due to their large range, migratory birds can either follow their climatic niche from one season to another (so-called ‘niche followers’ or ‘niche trackers’), or they experience varying climatic conditions. (i.e., ‘niche switchers’; Joseph and Stockwell 2000; Joseph 1996; Martínez-Meyer et al. 2004; Williams et al. 2017).

Pied cuckoo (*Clamator jacobinus*, Boddart, 1783) (Figure 1) is a type of crested cuckoo (Payne and Juana, 2020) found in the Indian and African Subcontinent and involved in inter and intracontinental migrations (Payne, 2005). This species is a brood parasite to different avifauna (Friedmann, 1964; Gaston, 1976; Johnsingh & Paramanandham, 1982; Payne, 2005) and a summer migrant to India (Friedmann, 1964; Whistler, 1928). There are three subspecies of this bird (Payne and Juana, 2020), viz. i. *C. jacobinus pica* (Hemprich and Ehrenberg, 1833) found in Southern Asia to Pakistan, northern India, Nepal, Tibet, Kashmir and Burma, and Africa south to Tanzania and Zambia (Gaston, 1976); ii. *C. jacobinus jacobinus* (Boddart, 1983); a resident subspecies found in southern India and Sri Lanka (Johnsingh & Paramanandham, 1982), and iii. *C. jacobinus serratus* (Sparrman, 1786); found in coastal South Africa; Zambezi River to the Cape region of South Africa. Pied Cuckoo is a summer migrant to India (Whistler, 1928; Friedmann, 1964) and also a brood parasite to several avifauna (Johnsingh and Paramanandham, 1982; Payne, 2005; Gaston, 1976; Friedmann, 1964). *C. jacobinus* parasitizes several host species (Friedmann, 1964), primarily babblers (genus *Turdoides*) in the plains and laughing-thrushes (genus *Garrulax*) in the hills (Friedmann, 1964; Gaston, 1976). In northern India, the subspecies *C. jacobinus. jacobinus* has been mentioned as resident (Payne, 2005), while the north Indian population is migratory (Ali and Ripley, 1983; Khachar, 1990).

**Figure 1.**
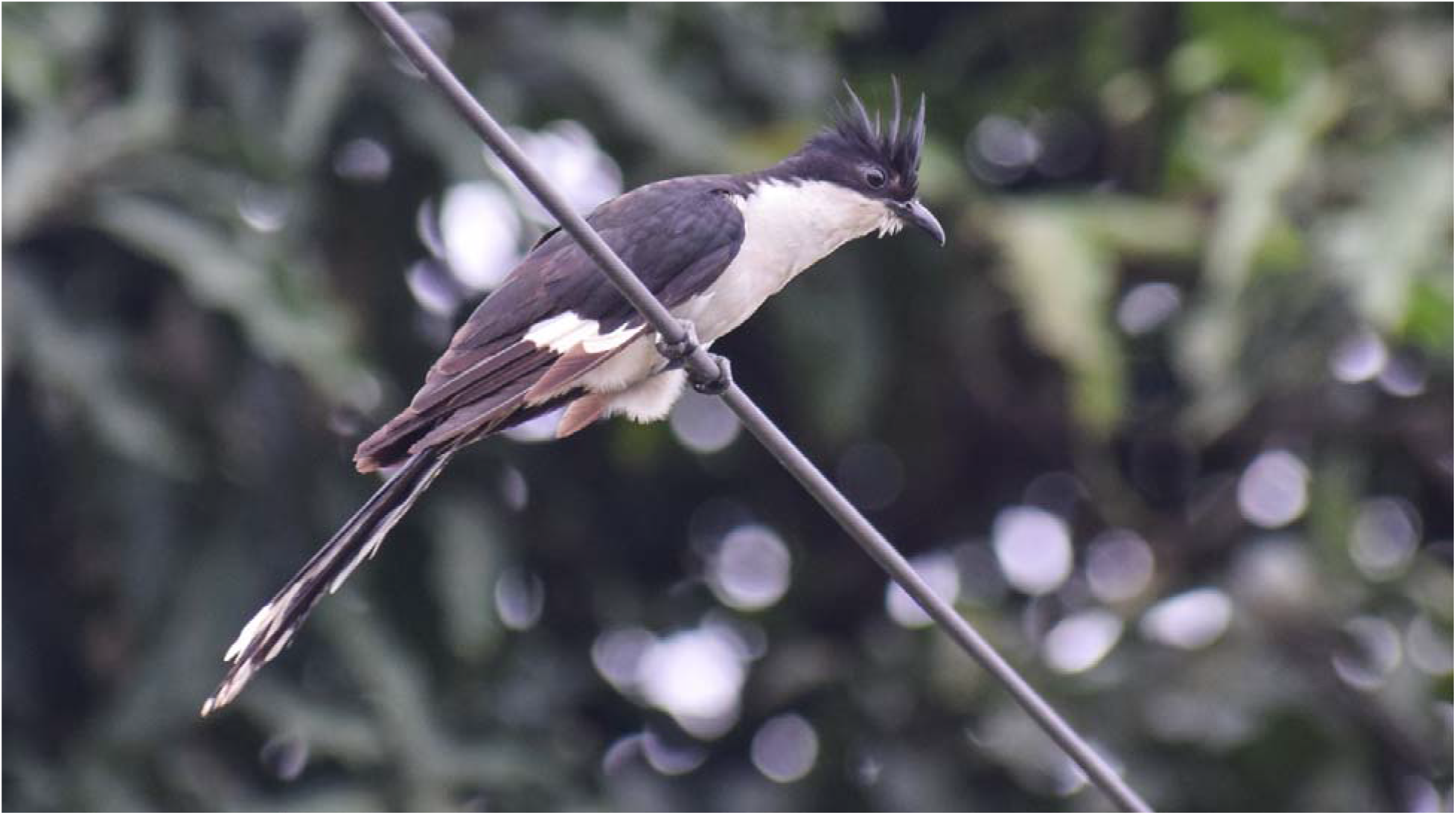
Photograph of a pied Cuckoo (*Clamator jacobinus*)

The species is deeply rooted in the regional culture and mythology of India. Historically, the advent of the monsoon for generations has been associated with the appearance of the Pied cuckoo in many parts of India (Ali and Ripley 1989), and the species is mentioned several times in ancient Indian literature and folklores (Kalidasa and Kale, 1974; Chopra, 2017). Ancient Hindu poetry refers to Pied cuckoos as *‘Chatak,’* who live on drops of rain (Abdulali 1972). In Kalidasa’s *Abhijñānaśākuntalam* the bird has been mentioned as “Divaukas,” who said to be present on Earth at certain times of the year (Chopra, 2017), denoting the migratory behaviour of the species. Nevertheless, unlike most migratory birds arriving in India that come from the northern hemisphere to winter, the Pied cuckoo is an exception in being a summer visitor to Northern India. It is known to make its appearance in several parts of Central and Northern India in the last week of May or early June, announcing the monsoon’s impending arrival with its unmistakably loud metallic call. India is a country with an agricultural economy; the monsoon is considered one of India’s most auspicious seasons. And so, the Pied cuckoo in North and Central India is a welcome sight. Reports indicate that the arrival of the Pied Cuckoo, coincides with the variation in the arrival of the monsoon winds (migrantwatch, 2013, (http://www.migrantwatch.in/blog/2013/04/04/does-the-pied-cuckoo-herald-the-monsoon/). A recent study by Madhusudan (2018) (https://birdcount.in/rain-bird-monsoon/) using citizen science data shows the correlation of the bird’s arrival in India with the southwest monsoons winds. Even though being a species with such ecology(Payne, 2005) and evolutionary history (Friedmann, 1984); apart from folk knowledge about the species, a few efforts to assemble relevant info in the early 1900s (Betts 1929; Pillai 1943; Simmons 1930; Whistler 1928), and few recent studies (Migrantwatch, 2013; Madhusudan, 2018), a very little information is available on its migration, monthly distribution pattern, its relation to the Indian monsoon and allied bioclimatic factors affecting its distribution in India.

Species Distribution Modelling (SDM) is used to predict species’ distribution across geographical space and time using species locations with a set of environmental data. It can help to fill knowledge gaps for poorly understood species by predicting suitable habitats within an area of interest, providing information about different habitat variables relevant to the species’ distribution, predicting responses to future climate conditions (Elith and Leathwick, 2009; Pearce and Boyce, 2006). SDM has varied practical applications ranging from conservation prioritization to effective habitat management of the species (Elith and Lithwick, 2009; Guisan and Thuiller, 2005; Williams, Willemoes, and Thorup, 2017;). Out of the many models used in SDM, Maxent (Maximum Entropy Species Distribution Modelling) (Phillips et al. 2006) is one of the leading algorithms for presence-only data in contemporary SDMs (Elith et al. 2006) and has outperformed other available presence only models (Anadón, Wiegand, and Giménez, 2012). There are numbers of studies have been done on predicting species migration through SDM’s using E-Bird, survey data, and satellite data (Coxen et al., 2017; Gschweng et al., 2012; Thorup et al., 2001; Revell and Somveille, 2017; Peterson, Ball and Cohoon, 2002; Pocewicz et al., 2013). E-Bird data is a high-quality data source for SDMs, and models based on E-Bird data can have high overlap in habitat suitability scores with models based on satellite tracking data (Coxen et al., 2017). The E-Bird data has also sometimes matched or exceeded performance compared to the systematically collected data (Valerie, Elphick and Tingley, 2019)

In this present study, we have used the Maximum entropy modeling approach (MaxEnt) (Philips et al., 2006) using data from EBird, and a set of environmental variables to predict the monthly distribution of *Clamator jacobinus* in India. This study aims to model the distribution pattern of *Clamator jacobinus* in different months and see the influence of environmental and climatic factors on its migration in the Indian Subcontinent.

## 2. Materials and Methods

### 2.1. Study Area

For modeling the species distribution, we have used entire India (20.5937° N, 78.9629° E) as our study area (Figure 2). India falls in the oriental biogeographic realm and has ten different biogeographic zones. The country’s total geographical area is 3,287,240 sq.km, divided into 28 states and 8 Union Territories. The climate of India is a majorly tropical monsoon type. However, it encompasses a varied range of climate conditions across a vast geographic scale and diverse topography. India experiences an annual mean temperature of 21 °C and an average rainfall of 1045mm. India has majorly four seasons, Winter (December-February), Summer (March-May), South-west monsoon season (June-September), Post monsoon season (October-November). The country’s climate is influenced by two seasonal winds – the southwest monsoon and the northeast monsoon. The north-east monsoon, generally known as winter monsoon, blows from land to sea. In contrast, the summer monsoon or the southwest monsoon blows from sea to land after crossing the Indian Ocean, the Arabian Sea, and the Bay of Bengal. The southwest monsoon carries most of the rainfall during a year in the country.

**Figure 2.**
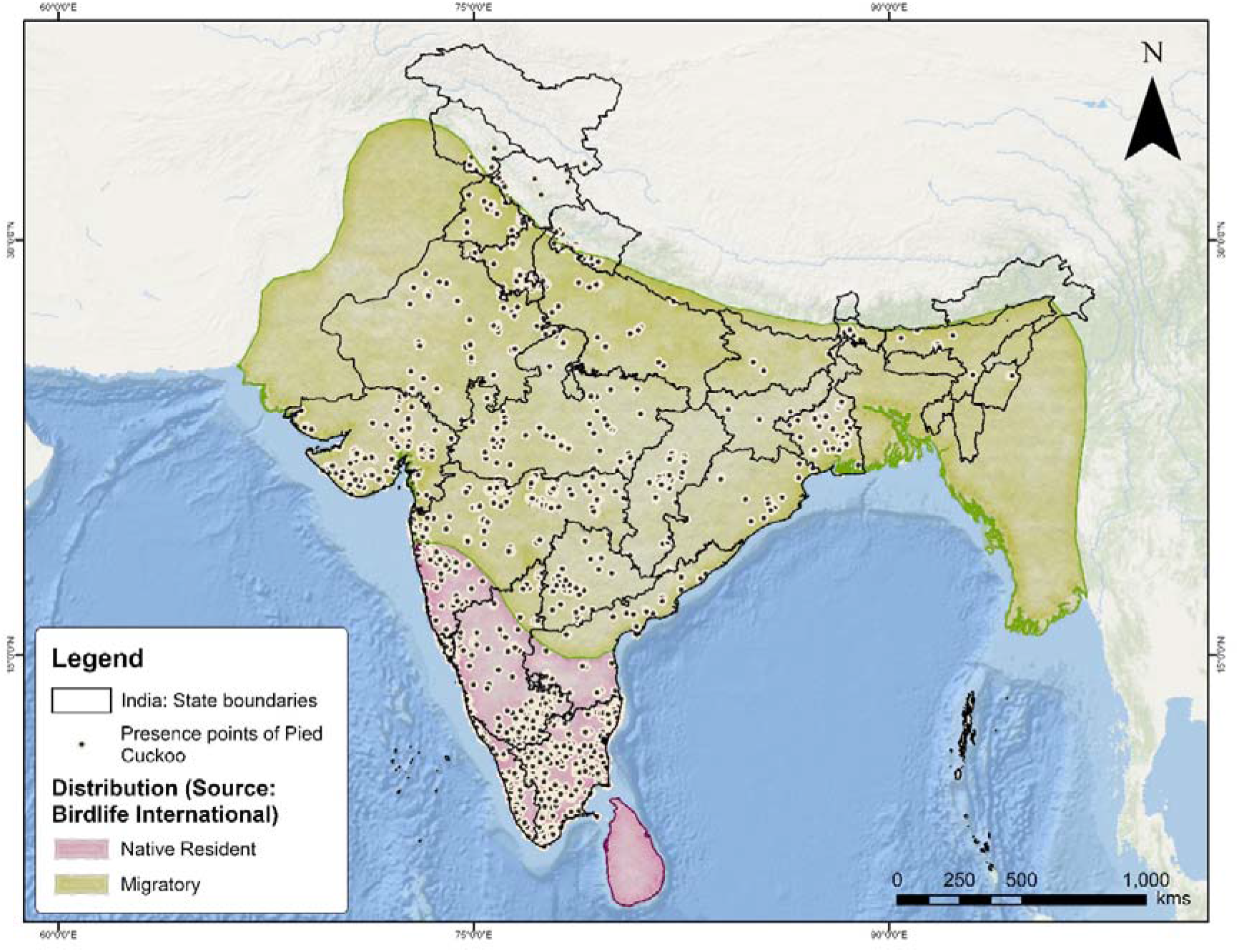
Study area for modeling the distribution of the Jacobin Cuckoo (*Clamator jacobinus*). Point locations denote the species presence (obtained from E-Bird). Native resident denotes resident species within the country whereas the migratory denotes the migratory population (Species range source: Birdlife International)

### 2.2. Species occurrence points

E-Bird (Sullivan et al., 2009) is a citizen science platform that exclusively focuses on distribution information about avifauna (Cox et al., 2012; Crall et al., 2010; Jackson et al., 2015; Lin et al., 2015). E-Bird records are submitted in checklist format listing the counts of each species encountered. Checklists include information on observation duration, distance traveled, and other method related metadata. We obtained these data by directly downloading the EBird Basic Dataset (https://ebird.org/science/download-ebirddata-products) on 30/1/2019.

Overall, out of 21294 downloaded occurrence points for the entire world, we clipped 8930 points for the year 2017-18 for India (Figure 2). However, citizen science data sometimes leads to spatial-clustering due to biased sampling (Zhang and Zhu, 2019). Thus, it is vital to remove the autocorrelated points (Boria et al., 2014; Fourcade et al., 2014). To minimize the error, the occurrence data were filtered and organized in sequential steps to improve quality. We rarefied occurrence data in ArcGIS using the Spatial Distribution Modeling toolbox (SDM toolbox v2.2.) (Brown, 2014) to spatially filter the data to a single point per 1 km^2^ grid and removed duplicate and spatially autocorrelated presence points. The filtered data was split for 12 months for running the models. We split the locations randomly in equal halves: one set of locations for calibration and another set for independent evaluation. The calibration data was split into two random subsets, training data (50%) and testing data (50%) (Supplementary figure. 1).

### 2.3. Bioclimatic variables

For modeling the distribution of *C. jacobinus*, we have used a set of environmental variables (Table 1) based on the available data, knowledge of species ecology, and potential factors affecting the species distribution. We have used WorldClim data V2.0 (Fick and Hijmans, 2017) for seven monthly bioclimatic variables, viz, Monthly minimum temperature (°C), maximum temperature (°C), average temperature (°C), Precipitation (mm), and water vapor pressure (kPa). Solar radiation (kJ m-2 day-1) and monthly Normalized Difference Vegetation Index (NDVI) (Didan, 2015) (Table 1). The Bioclimatic Data for each month were obtained from the WorldClim database version 2.0 (Fick and Hijmans, 2017, available at http://worldclim.org/version2) at 30-seconds (Approx. 1km2) resolutions. Monthly NDVI layers were extracted from the USGS Earth-Explorer database at a resolution of 250m. The geographic dimensions of all the downloaded layers were made similar (Pixel size 1 Km^2^) using the “SDM toolbox” (Brown, 2014) and “export to Circuitscape tool” by Jeff Jenness (http://www.jennessent.com/arcgis/Circuitscape_Exp.htm) in ArcGIS 10.7. Here we have not excluded any layers by correlation test as we intended to check the changing contribution of the bioclimatic variables with each month.

**Table 1.**
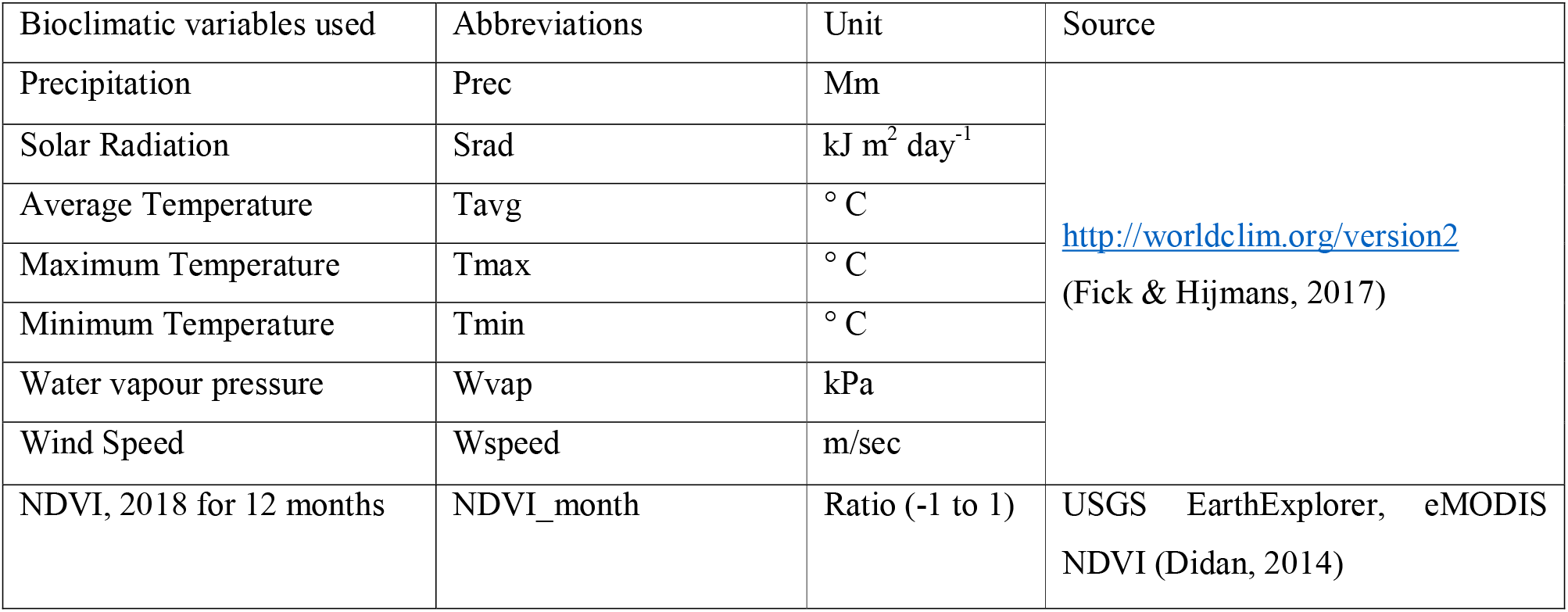
List of variables used for modeling the distribution.

### 2.4. Maximum entropy modeling

Maxent (Maximum Entropy Species Distribution Modelling) (Phillips et al. 2006) is one of the leading algorithms for presence-only data in contemporary SDMs (Elith et al. 2006) and has outperformed other available presence only models (Anadón et al. 2012; Kaliontzopoulou et al., 2008; and Wen et al. 2015; Wisz et al. 2008; Suárez-Seoane et al. 2008). We used MaxEnt 3.4.1 (Philips et al., 2006) in the R platform for estimating the monthly distribution pattern of Pied cuckoo in India. The calibration phase and choosing the best parameter are critical (Warren et al., 2010). For model calibration, model calibration and selection, final model creation, and evaluation, we have used ‘kuenm’ toolbox (Cobos, 2019) in R 3.5.0 (R Core Team, 2018).

#### 2.4.1. Model Calibration

For each month, we created 290 candidate models by combining ten values of regularization multiplier (0.5–5 at intervals of 0.1), and all 29 possible combinations of 5 feature classes (linear = l, quadratic = q, product = p, threshold = t, and hinge = h). We evaluated candidate model performance based on significance (partial ROC, with 500 iterations and 50 percent of data for bootstrapping), omission rates (E = 5%), and model complexity (AICc). Best models were selected according to the following criteria: (1) significant models with (2) omission rates ≤5%. Then, from among this model set, models with delta AICc values of ≤2 were chosen as final models.

#### 2.4.2. Final model building and evaluation

We created final monthly models for Pied cuckoo using the presence locations, selected model parameterizations, ten replicates by bootstrap with logistic output. A model’s application has little importance if the accuracy of the prediction is not measured (Pearson, 2010). We evaluated the model using Area under Receiver Operating Characteristics (ROC) in MaxEnt and a partial ROC (pROC) using a Kuenm package script. The ROC considers the sensitivity against the specificity of a model when new data is presented. The area under the Curve (AUC) of this ROC plot is a measure of the model’s overall accuracy. It ranges from 0 to 1, with 1 representing a model with perfect discrimination between sites where species are present and absent (Elith et al., 2006). Although widely used in the modeling literature, AUC’s effectiveness has been criticized as a way of predicting the accuracy of models (Lobo et al., 2008). As an alternative, the pROC has been proposed as an accurate test for model performance (Peterson et al., 2008). We present here both the AUC obtained from MaxEnt and the pROC results. The final map and AUC value are reported here for the average map of the ten replicates. Additionally, the ‘kuenm’ package was used to calculate the partial area under the ROC curve (pAUC) ratio to evaluate the performance of all models (Cobos et al. 2019). A pAUC ratio >1 indicates that the model has performed better than random chance. The pROC test was performed for each of the replicates and the average final map. Finally, we have used minimum training presence logistic threshold for creating binary maps (0 and 1) of the predicted distributions.

## 3. Results

### 3.1. Data filtering

We have used E-Bird data to model the monthly distribution of Pied cuckoo to predict its distribution throughout India in different months. We downloaded 21294 points downloaded initially for model building. After removing the duplicate and spatially clustered points total 3375 points were left for 2017-18. The data was split into twelve months for building the models (Supplementary Table 1).

### 3.2. Model Calibration and evaluation

We have initially calibrated 290 models for each month (total 3480 models) (Supplementary tables 2), among which 3453 number of models were statistically significant. Of these significant models, 562 (16%) met the omission criterion of 5% (Table 2). Finally, of the statistically significant, low-omission monthly models, models with the minimum AICc with selected regularization parameter values and features (Table 3) were used to create the final monthly models. The selection criteria values for each final model varied differently. Final models mean AUC ratio ranged from 1.116 to 1.824, omission rate at 5% ranging from 0.037 to 0.057 (Table 3).

**Table 2.**
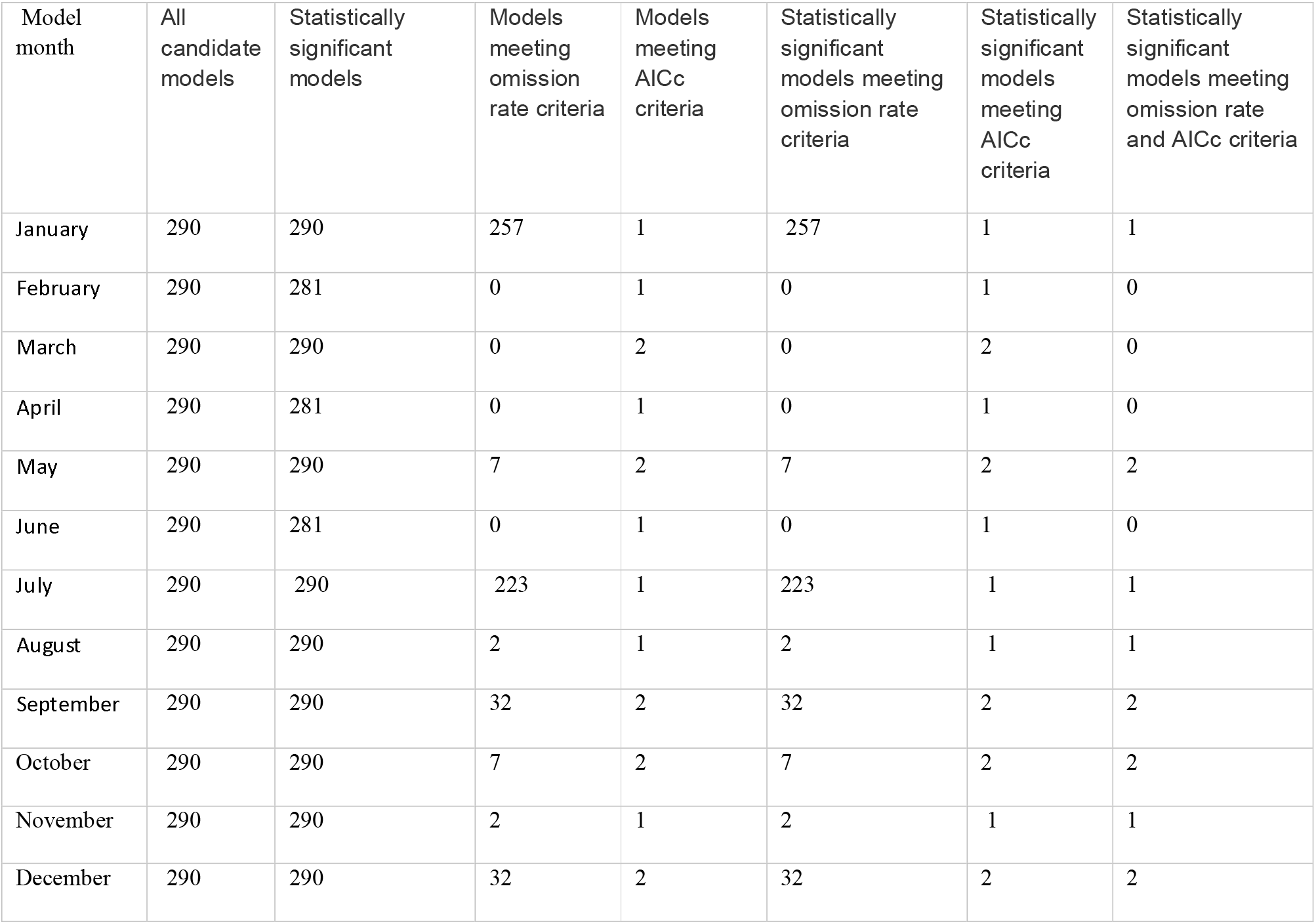
General Statistics of models that met distinct criteria

| Model month | All candidate models | Statistically significant models | Models meeting omission rate criteria | Models meeting AICc criteria | Statistically significant models meeting omission rate criteria | Statistically significant models meeting AICc criteria | Statistically significant models meeting omission rate and AICc criteria |
| --- | --- | --- | --- | --- | --- | --- | --- |
| January | 290 | 290 | 257 | 1 | 257 | 1 | 1 |
| February | 290 | 281 | 0 | 1 | 0 | 1 | 0 |
| March | 290 | 290 | 0 | 2 | 0 | 2 | 0 |
| April | 290 | 281 | 0 | 1 | 0 | 1 | 0 |
| May | 290 | 290 | 7 | 2 | 7 | 2 | 2 |
| June | 290 | 281 | 0 | 1 | 0 | 1 | 0 |
| July | 290 | 290 | 223 | 1 | 223 | 1 | 1 |
| August | 290 | 290 | 2 | 1 | 2 | 1 | 1 |
| September | 290 | 290 | 32 | 2 | 32 | 2 | 2 |
| October | 290 | 290 | 7 | 2 | 7 | 2 | 2 |
| November | 290 | 290 | 2 | 1 | 2 | 1 | 1 |
| December | 290 | 290 | 32 | 2 | 32 | 2 | 2 |

**Table 3.** Performance statistics for the best monthly models.

| Model month | Mean AUC ratio | Partial ROC | Omission rate at 5% | AICc | Delta AICc | W AICc | Num parameters |
| --- | --- | --- | --- | --- | --- | --- | --- |
| January | 1.76 | 0 | 0.057 | 6147.78 | 0 | 1 | 17 |
| February | 1.62 | 0 | 0.051 | 3248.176 | 0 | 1 | 25 |
| March | 1.474 | 0 | 0.051 | 3320.74 | 0 | 1 | 19 |
| April | 1.208 | 0 | 0.037 | 13436.16 | 0 | 0.987 | 69 |
| May | 1.36 | 0 | 0.043 | 4261.388 | 0 | 0.466 | 13 |
| June | 1.208 | 0 | 0.037 | 13436.16 | 0 | 0.987 | 69 |
| July | 1.231 | 0 | 0.046 | 12623.6 | 0 | 1 | 67 |
| August | 1.116 | 0 | 0.038 | 13046.17 | 0 | 1 | 26 |
| September | 1.263 | 0 | 0.05 | 11901.38 | 0 | 0.590 | 84 |
| October | 1.208 | 0 | 0.037 | 13436.16 | 0 | 0.987 | 69 |
| November | 1.231 | 0 | 0.046 | 12623.6 | 0 | 1 | 67 |
| December | 1.824 | 0 | 0.036 | 4985.472 | 0 | 1 | 38 |

### 3.3. Importance of bioclimatic variables

We have found the species is present throughout the year in southern India, and Northern Indian distribution is dynamic, highest in June (257400 Km^2^), and lowest in February (Total predicted area: 189638 km^2^). In Northern India, distribution started to increase from May before the onset of monsoon and after post-monsoon the distribution decreased gradually. The percent contribution of environmental variables differed between each monthly model (Figure 3). Water vapour pressure contributed highest before its arrival (highest 83% in April) and wind speed during the species departure (highest 42.8% in August). Precipitation was an important variable (24.8% contribution) in May, and NDVI contributed highest in July (26.4%).

**Figure 3.**
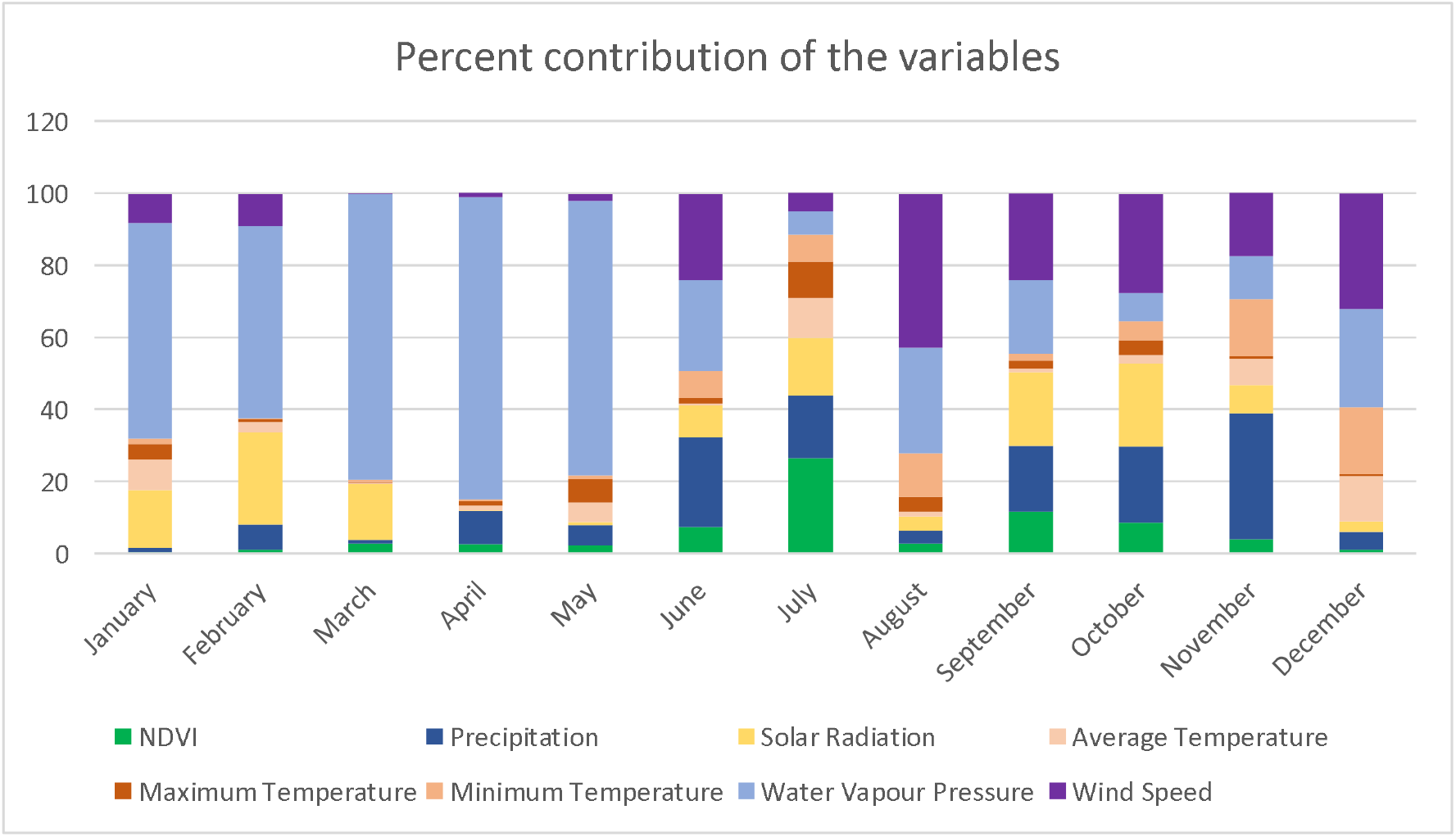
Percentage contribution of the bioclimatic variables used for monthly models of the Pied Cuckoo (*Clamator jacobinus*)

## 4. Discussion

In this study, we have used species occurrence points from e-bird and a set of environmental variables to model the monthly distributions of *C. jacobinus* in India. The monthly dynamic models produced in this study indicates the arrival of the bird to Northern India has a positive correlation with the onset of monsoon. The amount of contribution of the environmental variables varied every month (Figure 3)

### Winter distribution (December to February)

In these months, the species distribution (Figure 4) is restricted to Southern India (Majorly within Andhra Pradesh, Karnataka, Kerala, and Tamil Nadu). In these three months, temperature has been the primary contributing variable restricting the species distribution in Southern India, cumulatively contributing 46.4% in December. *C. jacobinus* is known to have a resident population in Southern India, and temperature is a critical factor restricting its distribution during the colder months of the year.

**Figure 4.**
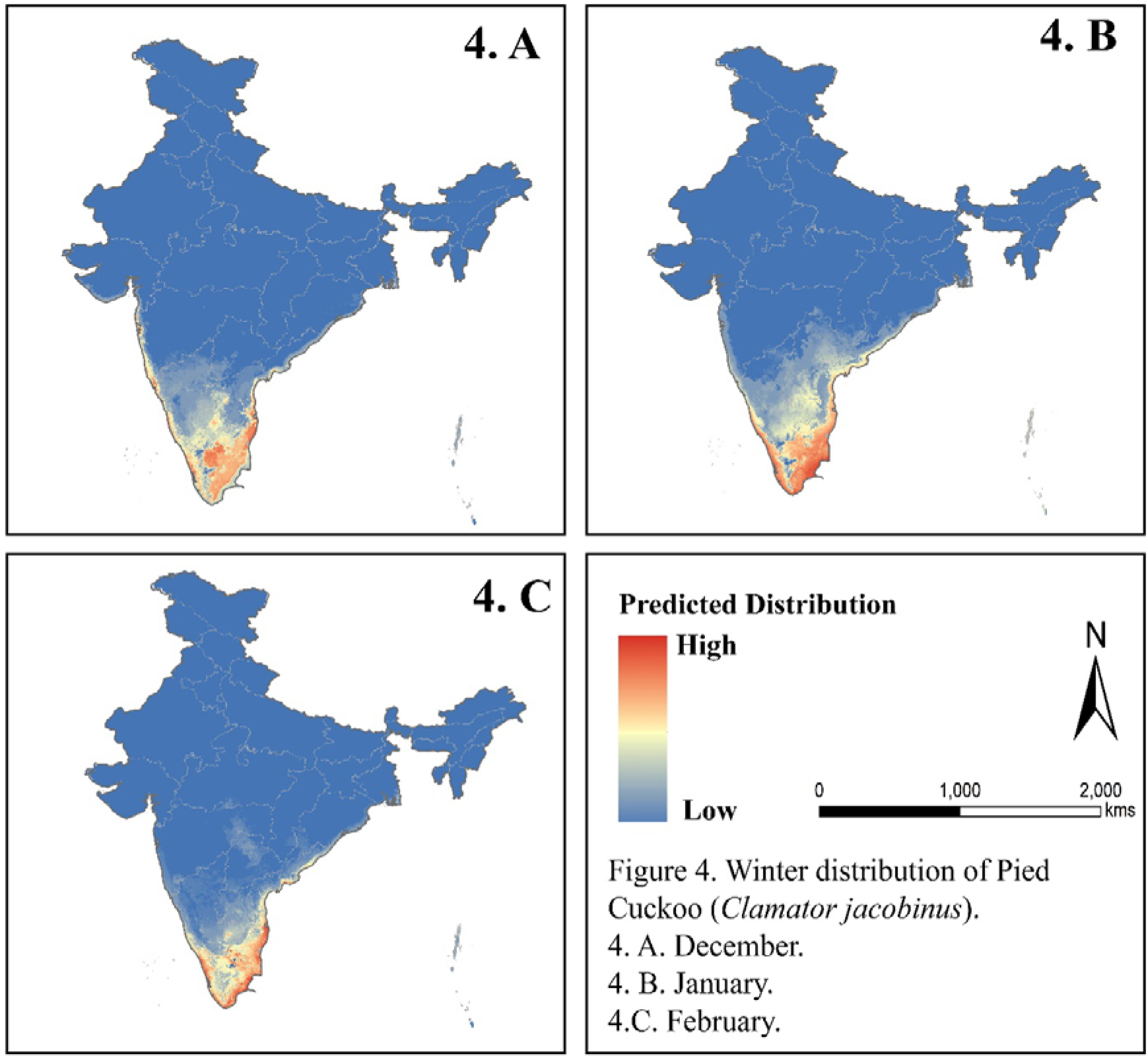
Winter distribution (4.A. December; 4.B. January, 4.C. February) of the Pied Cuckoo in India. Red colour denote higher value whereas blue is the lowest. The distribution of the species is limited in the Southern India in winter, lowest in February (4.C)

### Summer distribution (March to May)

Distributions in these months (Figure 5) shows that the species first enters the coasts of India, following a similar pattern like monsoon South-west monsoon winds, covering the coastal regions of at first and then gradually starts occupying the Northern and Central parts of India (There are anecdotal records of the species observed early in those regions compared to other areas). In May (Figure 5.C), the species reaches Northern India. Major variables contributing to the distribution for these months were water vapour pressure and wind speed (Figure 3).From March, when the species starts to arrive in the Northern Parts of India, the contribution of Water Vapour Pressure increased drastically (Figure 3); contributing as high as 83.8% in April and again starts decreasing (Lowest in July, contributing 6.5%) when the species reaches Northern India. Here, we hypothesize that the species is probably following the water vapour pressure. A high-water vapour value is a forerunner of rainfall, denoting species’ arrival in Northern India just before the monsoon.

**Figure 5.**
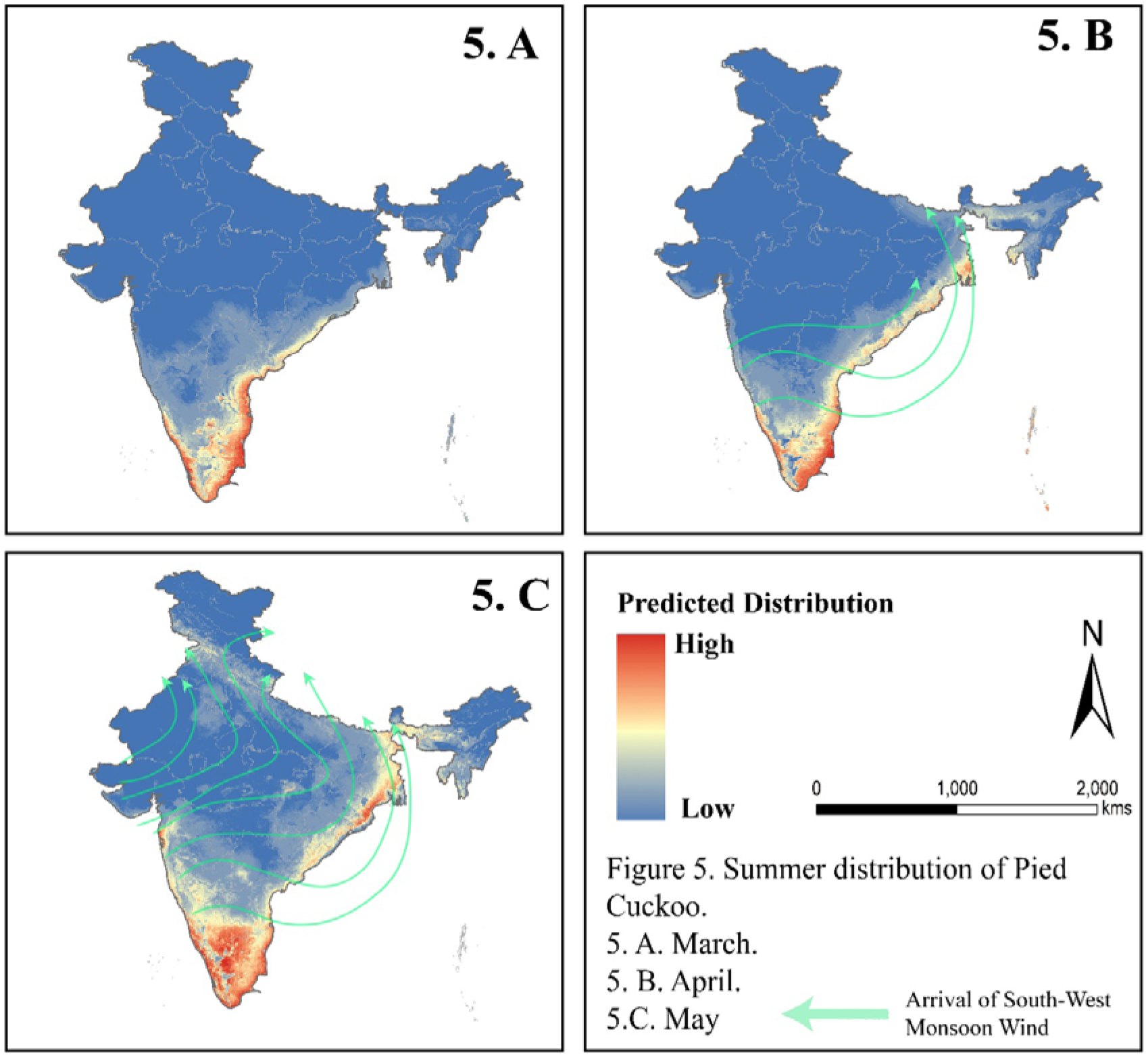
Predicted summer distribution (5.A. March; 5.B. April, 5.C. May) of the Pied Cuckoo (*Clamator jacobinus*) in India. Red colour denote higher value whereas blue is the lowest prediction.

### Monsoon and Post monsoon distribution (June-October)

In these months, monthly models predicted the species presence throughout Northern India, covering almost all the states of Northern India except the alpine regions. In June (Figure 5.A), Precipitation (24.8%) and Water Vapour Pressure (25.2%) have contributed maximum towards predicting the species distribution, representing the presence of Pied Cuckoo during the monsoon in Northern India. When monsoon arrives in India, the species is present the entire Northern plain parasitizing nests of different hosts (Friedmann, 1984) of respective regions. In July (Figure 6.B), however, NDVI contributes maximum (26.4%) (Figure 3) for predicting the species distribution. Cuckoos are specialist insectivores, specializing in caterpillar-based diets (Payne, 2005). These caterpillars (mainly *Eupterote modliflora* and *Macrobrochis gigas*) are abundant in rainy seasons when vegetation is green, and abundant food sources are available for the caterpillar. A recent study has found that cuckoos track high average ‘greenness’ (Thorup et al. 2017). Here, NDVI is considered a surrogate for the vegetation ‘greenness.’ By the August end, they parasitize its host’s nests and start preparing return migration. It is assumed that adults start their migration in October. Also, *C. jacobinus* juveniles start their return migration late without their parents, most likely in late October. In November (Figure 6.F), the species retreats from Northern India, leaving its resident population of Southern India (Majorly the states of Andhra Pradesh, Karnataka, Kerala, and Tamil Nadu). From August, the primary contributing variable is windspeed (Figure 3), factor aiding in its return migration.

**Figure 6.**
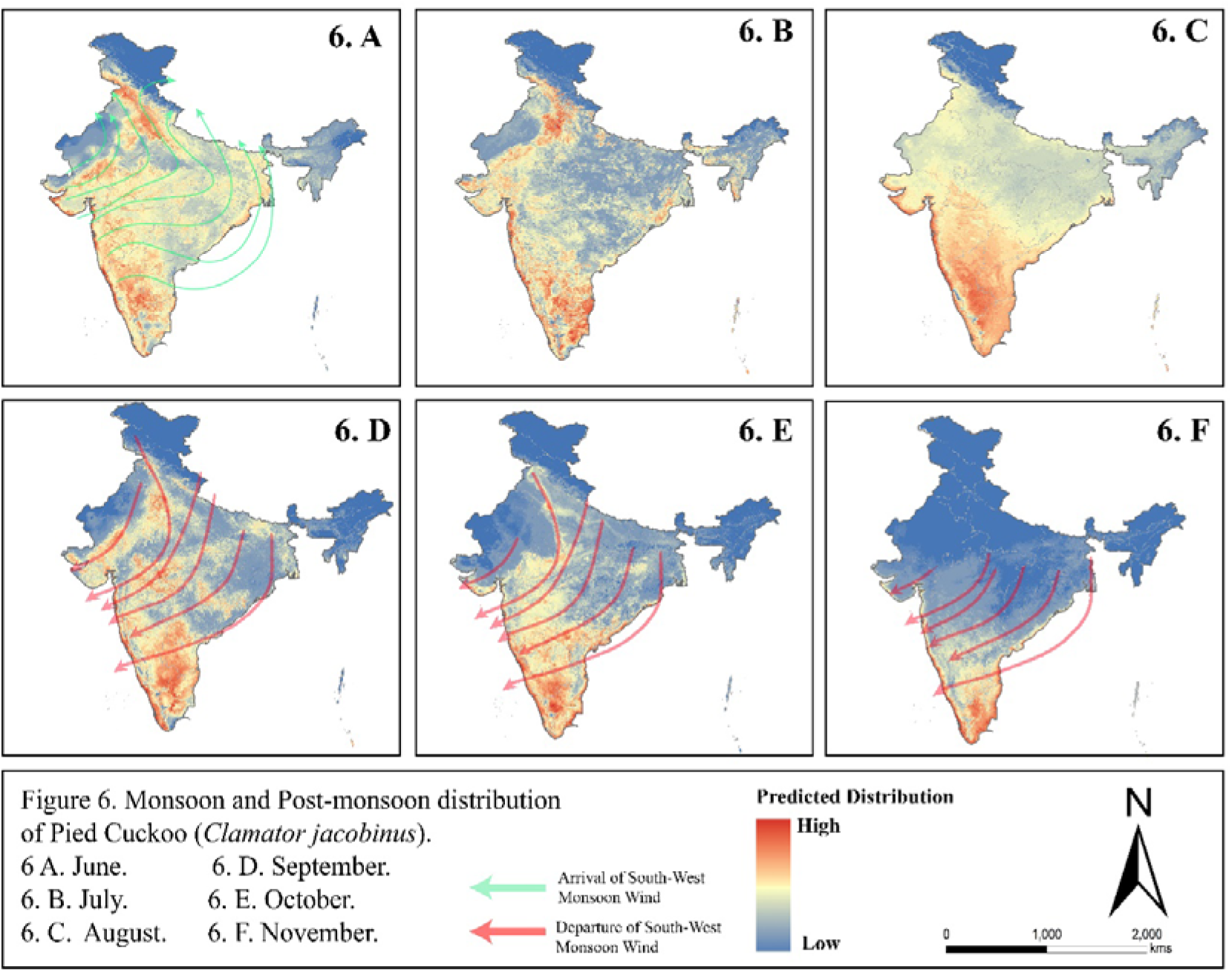
Monsoon and Post monsoon distribution (6. A June; 6. B. July; 6. C. August; 6. D. September; 6. E. October, 6. F. November; Red colour denote higher value whereas blue is the lowest prediction.

## 5. Conclusion

Here, we have modeled the monthly distribution of the species throughout India for the first time. A *latitudinal effect* could probably be why the species is present in the northern Part during summer but absent in winter. Many assumptions are present for its migration to date. It is assumed that the South Indian subspecies is resident and the northern India population is migratory. However, it could also be possible that the entire Indian population of this species is dynamic, where a part of the African population migrates and replaces the southern Indian population every year during summer, and the Southern India population migrates to Northern India for breeding. To verify these, we have tagged two Pied cuckoo individuals with 2gm satellite transmitters to identify its migration pathway and its population dynamics in the future.

SDM studies focusing on migratory species are infrequent. SDMs may greatly help define the spatial and temporal variation of habitat suitability for migratory avian species, but their use has been restricted to a few cases (Brambilla and Ficetola. 2012, Sardà-Palomera et al. 2012). However, new frameworks that incorporate such diverging species-environment relationships in time and space (Frans et al. 2018) may enable future studies. Ignoring these issues could, in turn induce an underestimation of areas needed for effective conservation (Runge et al. 2016) or an over-prediction of ranges in general (Reside et al. 2010). Hence, a careful selection of presence records is pivotal to limit such risks (Chamberlain et al. 2013). Our study has supported the hypothesis of migration of *Clamator jacobinus* and its link with the Indian monsoon’s arrival. Our approach provides a more concise understanding of monthly distributions of *C. jacobinus* throughout India, which helps understand the complex seasonal shifts in the distribution of such migratory birds.

**Figure 7.**
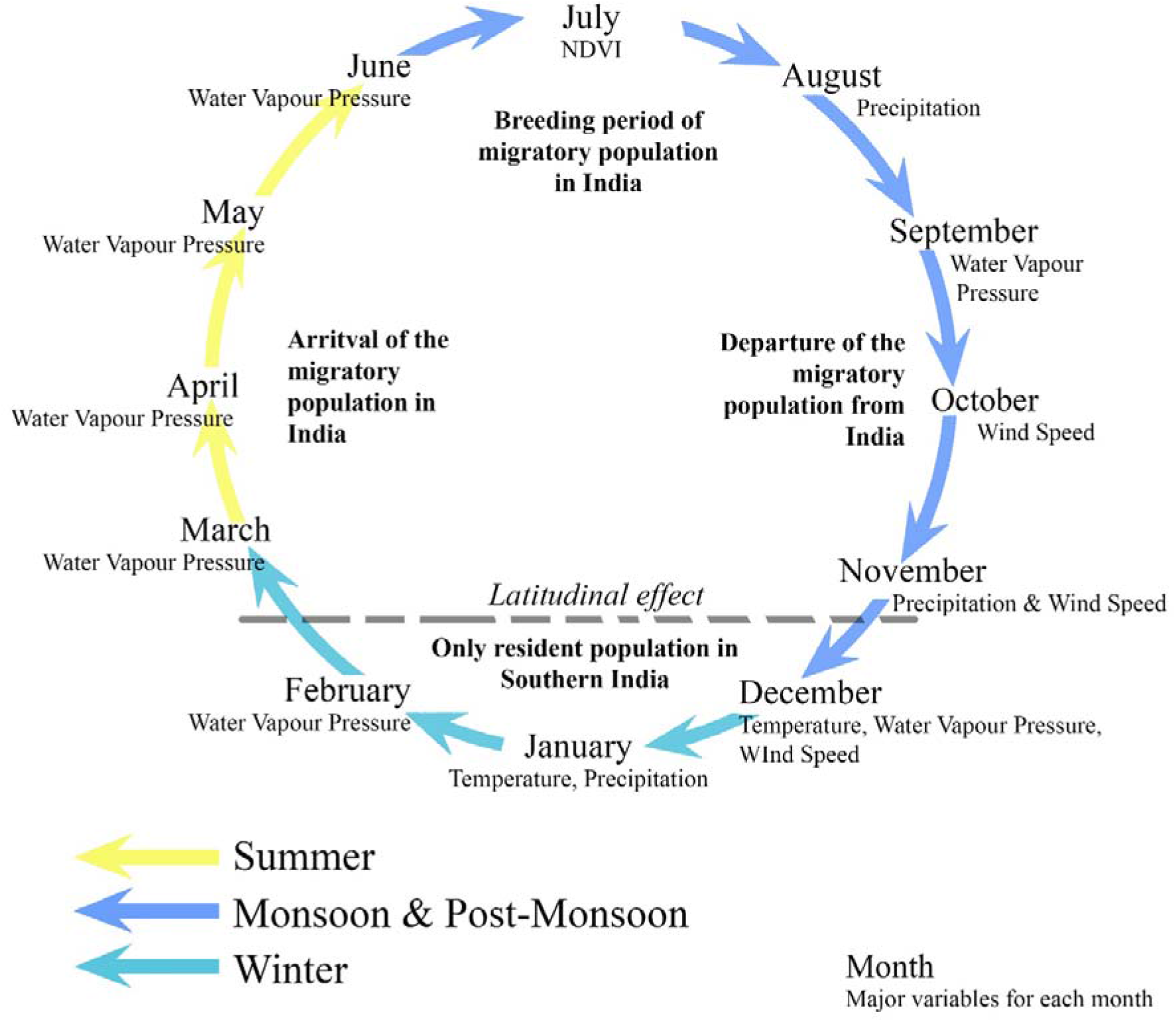
Summary of the migration cycle of Pied Cuckoo (*Clamator jacobinus*) in India and main bioclimatic variables affecting distribution in different months. More details of the contribution of each variable are provided in figure 3

## Supporting information

Supplementary figure 1 calibration results

Supplementary tables 2 performance statistics

Supplementary Table 1 location_information

## 6. Acknowledgments

The authors would like to thank Dean & Director, WII, for encouragement. We want to thank Anindita Debnath and Anukul Nath for their inputs. Funding for this paper was by the Department of Biotechnology (DBT), India, through “Indian Bioresource Information Network (IBIN) Geoportal Phase III: Enhancing BioResource Services, Institutional Linkages and Outreach (BT/Coord. II/01/04/2016)” project.

## 7. Author contributions

Debanjan Sarkar: Conceptualization, Data curation, Methodology, Analysis, Writing-Original draft preparation. Bharti Tomar: Data curation, Analysis. R. Suresh Kumar: Conceptualization, Supervision. Sameer Saran: Supervision. Gautam Talukdar: Conceptualization, Methodology, Investigation, Supervision, Writing-Original draft preparation.

