## Supplementary figures and images for "“Tracking the Rain Bird”: Modeling the monthly distribution of Pied cuckoo in India"

### Supplementary figure 1 calibration results

Supplementary figure 1. Calibration figures of the monthly models.

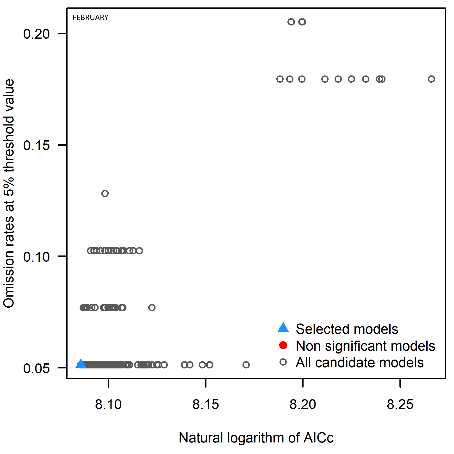

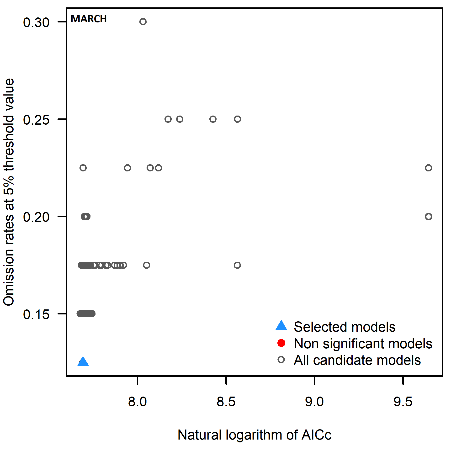

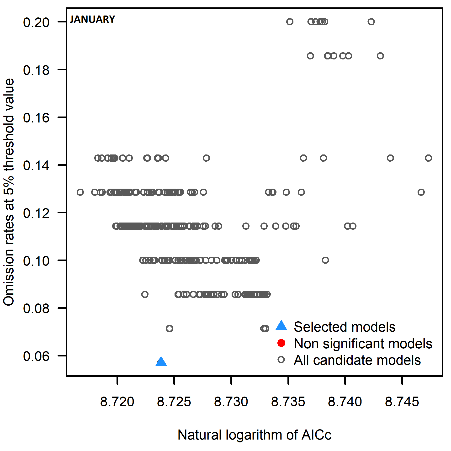

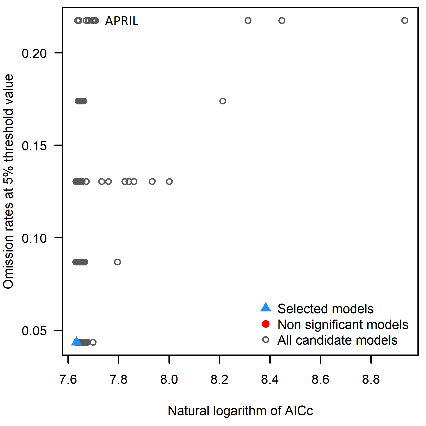

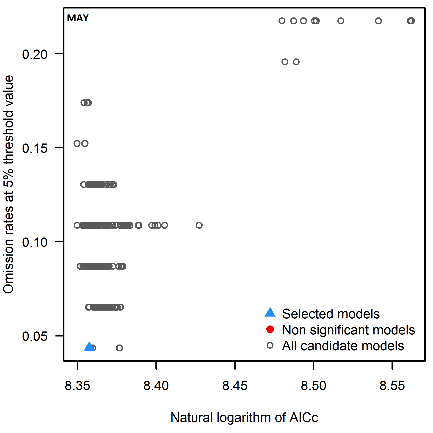

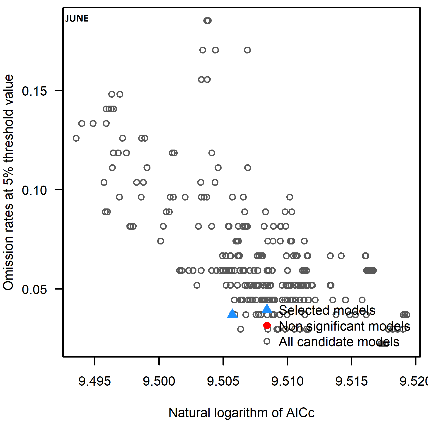

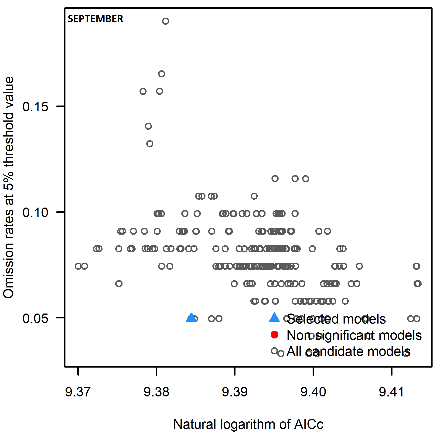

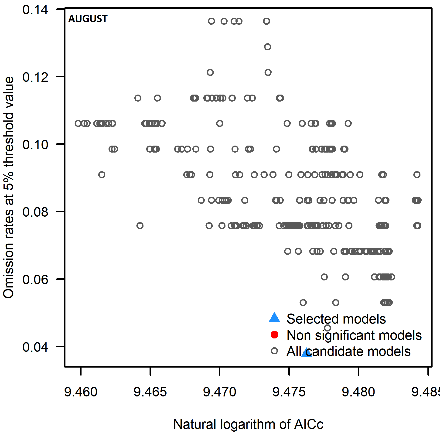

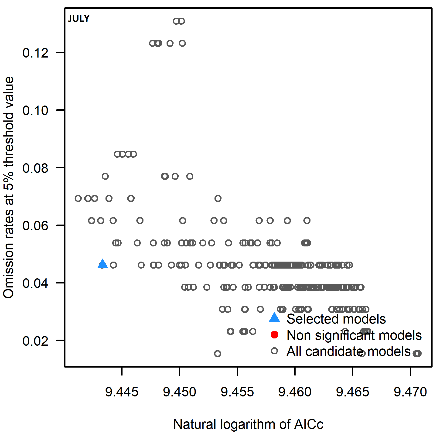

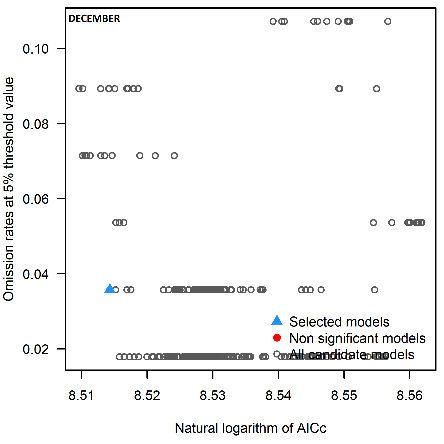

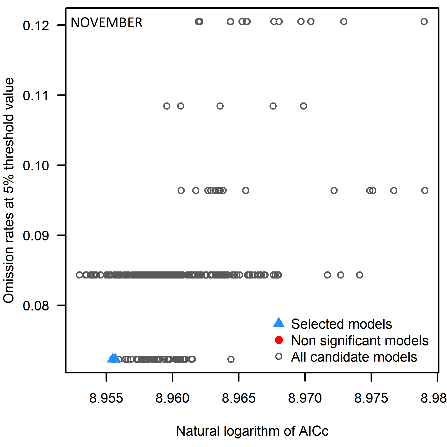

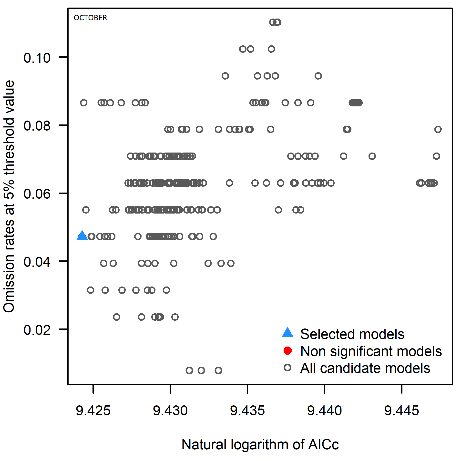
